# DPR-MEDIATED H_2_O_2_ RESISTANCE CONTRIBUTES TO STREPTOCOCCI SURVIVAL IN A CYSTIC FIBROSIS AIRWAY MODEL SYSTEM

**DOI:** 10.1101/2024.03.25.586644

**Authors:** Rendi R. Rogers, Christopher A. Kesthely, Fabrice Jean-Pierre, Bassam El Hafi, George A. O’Toole

## Abstract

The cystic fibrosis (CF) lung environment is conducive to the colonization of bacteria as polymicrobial biofilms, which are associated with poor clinical outcomes for persons with CF (pwCF). *Streptococcus* spp. is highly prevalent in the CF airway, but its role in the CF lung microbiome is poorly understood. Some studies have shown *Streptococcus* spp. to be associated with better clinical outcomes for pwCF, while others show that high abundance of *Streptococcus* spp. is correlated with exacerbations. Our lab previously reported a polymicrobial culture system consisting of four CF-relevant pathogens that can be used to study microbial behavior in a more clinically relevant setting. Here, we use this model system to identify genetic pathways that are important for *Streptococcus sanguinis* survival in the context of the polymicrobial community. We identified genes related to reactive oxygen species (ROS) as differentially expressed in *S. sanguinis* monoculture versus growth of this microbe in the mixed community. Genetic studies identified Dpr as important for *S. sanguinis* survival in the community. We show that Dpr, a DNA binding ferritin-like protein, and PerR, a peroxide-responsive transcriptional regulator of Dpr, are important for protecting *S. sanguinis* from phenazine-mediated toxicity in co-culture with *P. aeruginosa* and when exposed to ROS, both of which mimic the CF lung environment. Characterizing such interactions in a clinically relevant model system contributes to our understanding of microbial behavior in the context of polymicrobial biofilm infections.

**IMPORTANCE:** *Streptococcus* spp. is recognized as a highly prevalent pathogen in CF airway infections. However, its role in clinical outcomes for pwCF is poorly understood. Here, we leverage a polymicrobial community system previously developed by our group to model CF airway infections as a tool to investigate a *Pseudomonas-Streptococcus* interaction involving ROS. We show that protection against ROS is required for *S. sanguinis* survival in a clinically relevant polymicrobial system. Using this model system to study interspecies interactions contributes to our broader understanding of the complex role of *Streptococcus* spp. in the CF lung.

## INTRODUCTION

The cystic fibrosis (CF) airway is characterized by the presence of thick, dehydrated mucus that impairs mucociliary clearance and creates an environment conducive to bacterial colonization. Persons with CF (pwCF) commonly develop chronic lung infections that microbiome studies have shown to be diverse and polymicrobial (1–3). Although streptococci are now recognized as a central component of the CF lung microbiome (4–7), their specific role in CF lung disease remains unclear. Previous findings have shown that an increased relative abundance of streptococci in the lung, including those originating from the oral microflora, corresponds to less severe impacts on lung disease compared to infection with other CF-associated microbes (5, 6, 8–10). In contrast, other studies have shown that a higher abundance of streptococci is more likely in pwCF upon receiving treatment for severe airway symptoms (11, 12). The *Streptococcus anginosus* group (SAG, also known as the *Streptococcus milleri* group) are specifically correlated with increased exacerbation when they are the predominant species in the lung (4, 13–15). Together, these data underscore the complex and multifaceted role of streptococci in the context of the CF airway (16).

Previous findings indicate that streptococci in the CF lung may interact with other CF pathogens in ways that could alter bacterial physiology and virulence (2, 10, 17–24). Notably, coculture experiments have suggested that *Streptococcus* spp. engage in both positive and negative interactions with the prominent CF pathogen *Pseudomonas aeruginosa*. For example, *P. aeruginosa* can positively impact *Streptococcus* spp. biofilm formation via secretion of exopolysaccharides (19, 24). Conversely, *P. aeruginosa* can negatively affect *Streptococcus* spp. growth via rhamnolipid secretion (20). Furthermore, *Streptococcus* spp. have been shown to enhance virulence factor production by *P. aeruginosa* (21, 23), but streptococci can also inhibit *P. aeruginosa* growth and viability via production of reactive oxygen species (ROS) and reactive nitrogen species (RNS) (17, 18, 22). It is important to note nearly every study cited here was performed in different media and under growth conditions that do not necessarily reflect the CF lung environment, thus it is difficult to determine which of these interactions might be clinically relevant.

Work from our lab has aimed to address existing gaps in our understanding of polymicrobial biofilms within the framework of CF infections by using a novel, clinically-informed model community consisting of four CF-relevant microbes (25). The model community consists of *P. aeruginosa, Staphylococcus aureus, Streptococcus sanguinis,* and *Prevotella melaninogenica* cultured in artificial sputum medium (ASM) supplemented with mucin and grown under anoxic conditions, simulating the mucus plug environment where microbes are often found in the CF lung (26, 27).

Using the aforementioned co-culture system, we have reported that several tested streptococci exhibit increased growth when in the presence of other microbial partners (25). Here, in an attempt to understand the community-specific genes important for the establishment of streptococci in a polymicrobial environment, we leverage RNA-Seq data from a recent study involving this polymicrobial community (28) to identify *S. sanguinis* genes that are differentially expressed when *S. sanguinis* is grown in coculture compared to monoculture. Multiple genes identified as highly downregulated in coculture are involved in the ROS response. We show that one of these genes, the DNA-binding ferritin-like protein Dpr, as well as a Fur-family transcriptional regulator, PerR, a known regulator of Dpr in other streptococci, are important for protecting *S. sanguinis* in coculture with *P. aeruginosa*, likely via mitigating phenazine-and H_2_O_2_-mediated toxicity. These findings suggest that a robust ROS response is a potential key factor for *S. sanguinis* survival in the CF airway, particularly in the context of a polymicrobial community.

## RESULTS

### *S. sanguinis* genes required for the ROS response are differentially expressed in monoculture compared to co-culture

We previously performed an RNA-Seq analysis of a CF-relevant polymicrobial community consisting of *P. aeruginosa* PA14*, S. aureus* Newman*, S. sanguinis* SK36, and *P. melaninogenica* ATCC 25845 (28). This analysis showed that nearly five hundred *S. sanguinis* genes are significantly differentially expressed when grown in the mixed community compared to growth as a monoculture. To investigate potential genetic pathways that contribute to *Streptococcus* microbe-microbe interactions in the CF airway, we identified those genes most differentially expressed in the mixed community compared to the monoculture (**Figure S1, Supplemental Table S1**). Our analysis found five genes involved in *S. sanguinis* response to ROS that are significantly and robustly upregulated in the monoculture when compared to the mixed community (**Figure 1A**).

**Figure 1.**
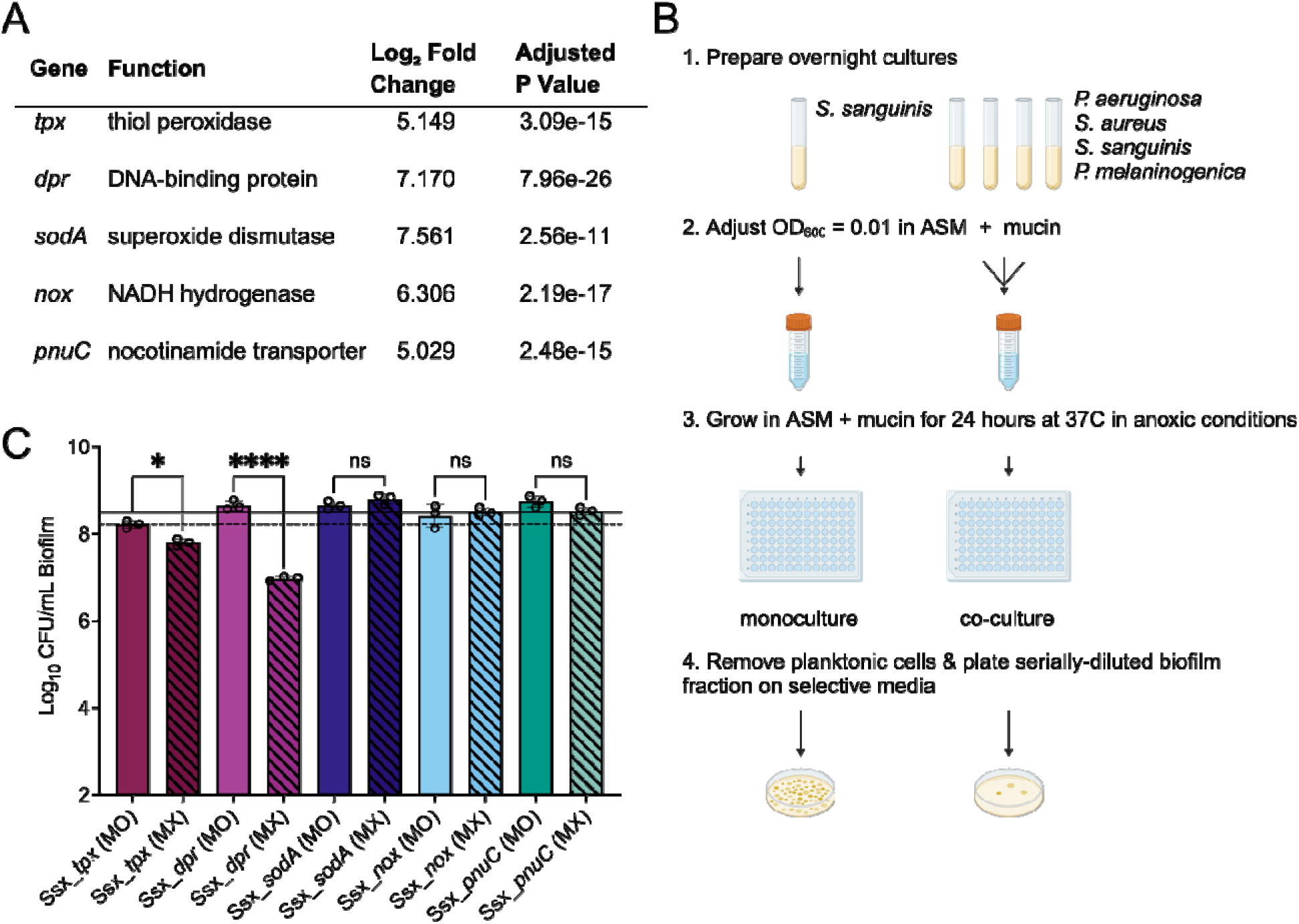
*S. sanguinis* ROS-related genes are differentially expressed in monoculture versus co-culture, and contribute to *S. sanguinis* survival in a mixed community. (**A**) Name and function of genes involved in oxidative stress response that are differentially expressed in monoculture versus the mixed community along with log_2_ fold change in the monoculture compared to the four-species mixed community. These genes are all significantly more highly expressed in monoculture versus co-culture. The adjusted P value using a Bonferroni correction is shown in the right-most column. (**B**) Experimental outline of the microbial growth assay used throughout this study. (**C**) Colony forming unit (CFU) counts for mutants grown in monoculture and the mixed community for 24 hrs. MO = monoculture, MX = mixed community. Black lines show wild-type (WT) *S. sanguinis* CFU counts across all experiments. Solid line = monoculture average. Dashed line = mixed community average. CFUs were performed by plating on medium selective for the growth of *Streptococcus* spp. Each data point presented in a column represents the average from at least three technical replicates performed at least on three different days (n=3). Statistical analysis was performed using ordinary one-way ANOVA and Tukey’s multiple comparisons posttest with *, p<0.05; ****, p<0.0001, ns = non-significant. Error bars

Next, we tested the impact of each gene shown in **Figure 1A** on *S. sanguinis* viability when grown in the mixed community as compared to the monoculture. Mutants were obtained from a comprehensive SK36 mutant library (29) and grown in a monoculture and mixed culture as previously described (**Figure 1B**) (25). Of the five mutants tested, only two showed a significant reduction in viability in the mixed culture compared with the Ssx_*tpx* mutant strain showing a 0.411 log_10_ reduction in biofilm viable counts and the Ssx_*dpr* mutant showing a 1.675 log_10_ reduction in biofilm growth quantified by CFUs compared to the WT (**Figure 1C**, **Figure S2**). Surprising, despite the higher expression of these genes in monoculture, none of the mutants tested showed a significant reduction in viability compared to the WT in monoculture (**Figure 1C**). Because the Ssx_*dpr* mutant showed the greatest reduction in viability, we narrowed the focus of this study to investigate Dpr.

### Loss of PerR function results in poor *S. sanguinis* growth in the mixed community

It has been previously shown that PerR functions as a peroxide responsive transcriptional regulator and is involved in regulating *dpr* gene expression as well as other genes required for the ROS response in *Streptococcus* spp. (30–33). Therefore, we hypothesized that the observed reduction in viability of the Ssx_*dpr* mutant in the mixed community may be dependent on PerR regulation. First, we conducted viability assays as described in the previous section (**Figure 1B**) with a Ssx_*perR* mutant in parallel with the Ssx_*dpr* mutant (**Figure 2A**) and observed that the Ssx_*perR* mutant mirrors the phenotype of the Ssx_*dpr* mutant. These findings indicated that PerR may positively regulate the *dpr* gene in *S. sanguinis*. Complementation of the Ssx_*dpr* or Ssx_*perR* mutant restored growth in the mixed community to a level not statistically significantly different from the growth of these mutants in monoculture (**Figure 2A**).

**Figure 2.**
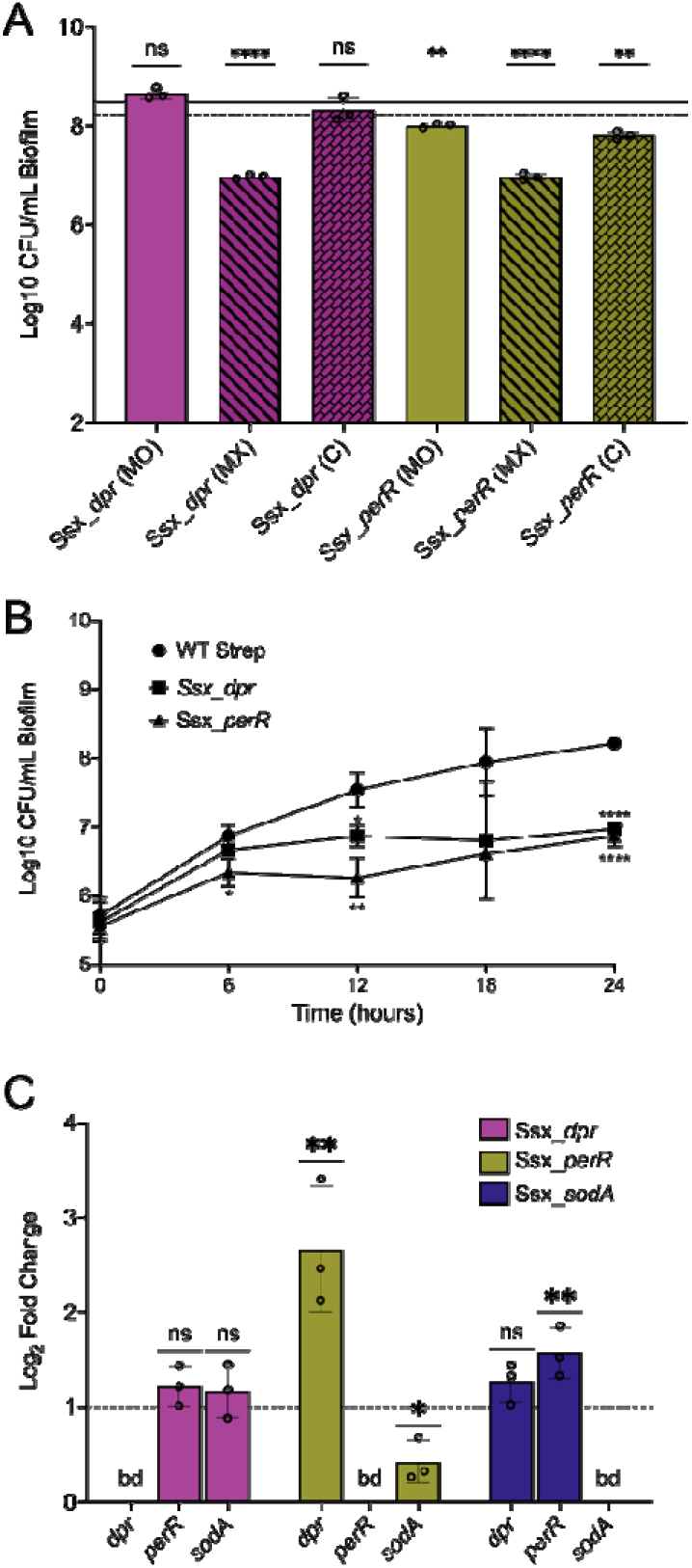
Loss of PerR, a *dpr* repressor, results in poor *S. sanguinis* growth in the mixed community. (**A**) CFU counts for mutants and their complements grown in monoculture and the mixed community. MO = monoculture, MX = mixed community, C = complemented in the mixed community. Black lines show wild-type (WT) *S. sanguinis* CFU counts across all experiments. Solid line = monoculture average. Dashed line = mixed community average. CFUs were performed by plating on medium selective for the growth *Streptococcus* spp. Each data point represents the average from at least three technical replicates performed at least on three different days (n = 3). Statistical analysis was performed using ordinary one-way ANOVA and Tukey’s multiple comparisons posttest with *, p<0.05; ****, p<0.0001, ns = non-significant. Statistics shown here compare mutant growth to WT growth in the same condition (MO to MO, MX to MX, MX to C). Error bars represent SD. CFU counts for other bacteria in the mixed community are shown in **Supplemental Figure S3**. (**B**) CFU counts for mutants measured every six hours over the course of 24 hours. CFUs were performed by plating on medium selective for the growth of *Streptococcus* spp. Each data point represents the average from at least three technical replicates performed at least on three different days (n = 3). Statistical analysis was performed using ordinary one-way ANOVA and Tukey’s multiple comparisons posttest with *, p<0.05; **, p<0.01; ****, p<0.0001, no notation = non-significant. Error bars represent SD. (**C**) Log_2_ fold change in expression of *dpr, perR,* and *sodA* genes in the Ssx_*dpr,* Ssx_*perR,* and Ssx_*sodA* mutant backgrounds compared to the WT, which is represented by the dashed line. Change in gene expression was calculated using the ΔΔ*C*_T_ method. Each data point represents the average from at least three technical replicates performed at least on three different days (n = 3). Statistical analysis was performed using ordinary one-way ANOVA and Tukey’s multiple comparisons posttest with *, p<0.05; ****, p<0.0001, ns = non-significant. Error bars represent SD.

Next, to test the pattern of growth of these mutants in the mixed community over time, we conducted the same viability assay (**Figure 2A**), this time quantifying CFUs at 6, 12, 18, and 24 hours post-inoculation (**Figure 2B**). While wild-type (WT) *S. sanguinis* grows throughout the 24-hour incubation period, we observed that both the Ssx_*dpr* and Ssx_*perR* mutants grow at a comparable rate to the WT for the first six hours, after which their growth plateaus. These data further support the hypothesis that loss of function of PerR and Dpr is responsible for the observed reduction in viable bacterial counts.

### PerR negatively regulates *dpr* gene expression

To test the hypothesis that PerR is a positive regulator of *dpr*, we quantified expression of *dpr* and another known PerR-regulated gene, *sodA*, in both the WT and Ssx_*perR* backgrounds using qPCR. We observed that *dpr* expression is significantly increased in the Ssx_*perR* background, while *sodA* expression, as expected (31), is significantly decreased (**Figure 2C**). These data indicate that PerR functions to repress the transcription of *dpr* in *S. sanguinis*. Furthermore, this result suggests that the observed loss of growth phenotypes of the Ssx_*dpr* and Ssx_*perR* mutants in the community may result from different mechanisms. We address this idea further in the Discussion. We also note that the Ssx_*dpr* mutant showed no change in *sodA* or *perR* gene expression, while mutating *sodA* does result in a modest by significant increase in *perR* gene expression.

### *P. aeruginosa* PA14 secreted phenazines cause poor growth for the *perR* and *dpr* mutants

To investigate the mechanism by which incubation in the polymicrobial community is causing the observed defect in the growth of the Ssx_*dpr* and Ssx_*perR* mutants, we conducted pairwise co-culture assays and found that growing the Ssx_*dpr* and Ssx_*perR* mutants with WT *P. aeruginosa* alone is sufficient to reproduce the phenotype observed in the four-species mixed community (**Figure S4**).

Next, we cultured the Ssx_*dpr* and Ssx_*perR* mutants in the presence of sterilized, cell-free supernatant from WT *P. aeruginosa* to test if secreted factors cause reduced growth for the Ssx_*dpr* and Ssx_*perR* mutants. The Ssx_*dpr* mutant shows a significant reduction in CFU counts in the presence of *P. aeruginosa* supernatant, although the defect is lesser than what is observed in the presence of live *P. aeruginosa* (**Figure 3A**). Interestingly, the Ssx_*perR* mutant also showed a reduction in CFU counts, although this reduction was not statistically significant (**Figure 3A**). These data suggest that a secreted factor might account for, at least in part, the reduced viability of the Ssx_*dpr* and Ssx_*perR* mutants. Based on these data, we sought to investigate which *P. aeruginosa* secreted factors might be responsible for the observed growth defects of these mutants in the community.

**Figure 3.**
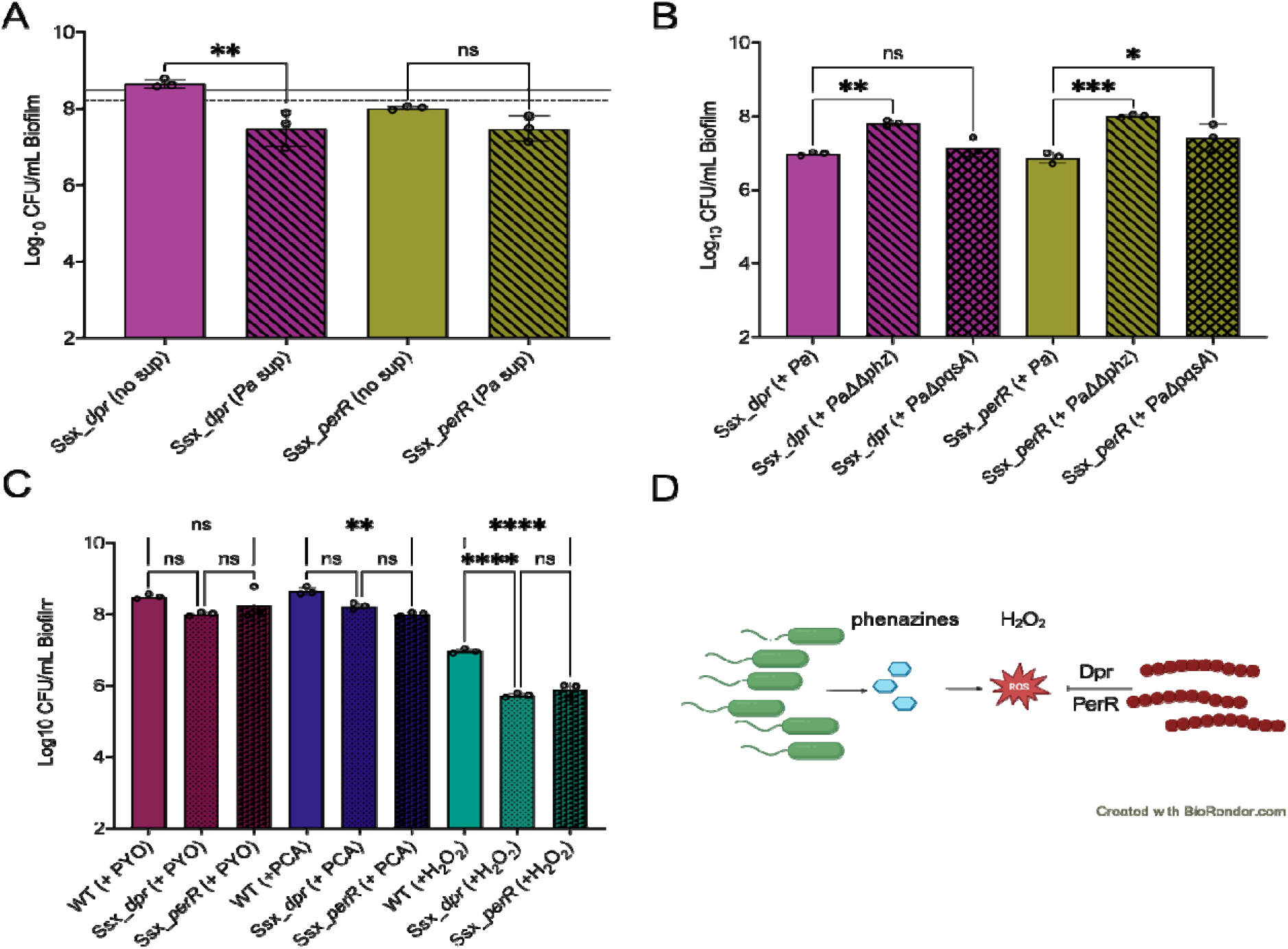
*P. aeruginosa* secreted phenazines cause poor growth in Ssx_*perR* and Ssx_*dpr* mutants. (**A**) CFU counts for *S. sanguinis* mutants grown in ASM supplemented with *P. aeruginosa* supernatant (*Pa* sup) or with a non-supernatant control (no sup). Black lines show wild-type (WT) *S. sanguinis* CFU counts across all experiments. Solid line = monoculture average. Dashed line = mixed community average. CFUs were performed by plating on medium selective for the growth of *Streptococcus* spp. Each data point represents the average from at least three technical replicates performed at least on three different days (n = 3). Statistical analysis was performed using ordinary one-way ANOVA and Tukey’s multiple comparisons posttest with *, p<0.05; ****, p<0.0001, ns = non-significant. Error bars represent SD. (**B**) CFU counts for *S. sanguinis* mutants grown in co-culture with WT *P. aeruginosa* (+ *Pa*), a *P. aeruginosa* phenazine mutant (*+ Pa*ΔΔ*phz*), or a *P. aeruginosa* quinolone signal mutant (+ *Pa*ΔΔ*pqs*). CFUs were performed by plating on medium selective for the growth *Streptococcus* spp. Each data point represents the average from at least three technical replicates performed at least on three different days (n = 3). Statistical analysis was performed using ordinary one-way ANOVA and Tukey’s multiple comparisons posttest with *, p<0.05; ****, p<0.0001, ns = non-significant. Error bars represent SD. CFU counts for all *S. sanguinis,* and *P. aeruginosa* strains tested in co-culture are shown in **Supplemental Figure S5**. (**C**) CFU counts for *S. sanguinis* WT and mutants grown in ASM supplemented with 50 uM pyocyanin (+ PYO), 50 uM phenazine-1-carboxylic acid (+ PCA), or 0.1% H_2_O_2_. CFUs were performed by plating on medium selective for the growth *Streptococcus* spp. Each data point represents the average from at least three technical replicates performed at least on three different days (n = 3). Statistical analysis was performed using ordinary one-way ANOVA and Tukey’s multiple comparisons posttest with *, p<0.05; ****, p<0.0001, ns = non-significant. Error bars represent SD. (**D**) A model demonstrating a *P. aeruginosa*-*S. sanguinis* interaction. In this study, we show evidence supporting the hypothesis that Dpr and PerR are important in protecting *S. sanguinis* from H_2_O_2_ formed as a downstream product of *P. aeruginosa* secreted phenazines.

Previous work showed that *P. aeruginosa* phenazine production is correlated with increased production of H_2_O_2_ (34). Based on this literature, we hypothesized that disrupting phenazine production by *P. aeruginosa* would rescue the growth defect observed in the Ssx_*dpr* and Ssx_*perR* mutants in co-culture. We also chose to test other factors known to be secreted by *P. aeruginosa*, particularly those with known roles in microbe-microbe interactions and virulence factors, including the *Pseudomonas* quinolone signal (PQS) (35, 36). Of the nine mutants tested (**Figure S5**), the *P. aeruginosa* PA14 Δ*phzA1-G1,A2-G2* mutant (37), which lacks the ability to produce phenazines, and the *P. aeruginosa* PA14 Δ*pqsA* (38), which lacks the ability to produce PQS, were the only mutants that conferred a significant increase in viability for the Ssx_*dpr* or Ssx_*perR* mutants (**Figure S5**). The data show that eliminating production of phenazines, but not PQS, rescues the reduced viability phenotype for both Ssx_*dpr* and Ssx_*perR* mutants (**Figure 3B**).

We next tested the effects of exogenous phenazine molecules pyocyanin (PYO) and phenazine-1-carboxylic acid (PCA) as well as hydrogen peroxide (H_2_O_2_) under our standard model conditions. We found that PYO does not cause a significant viability defect for the Ssx_*dpr* and Ssx_*perR* mutants compared to the wild type (WT), while PCA causes a significant viability defect in the Ssx_*perR* mutant and a small, but non-significant reduction for the Ssx_*dpr* mutant compared to the WT. Peroxides, which can be generated by phenazines (34) causes a significant viability defect for both Ssx_*dpr* and Ssx_*perR* mutants compared to the WT (**Figure 3C**). These data taken together, along with previous studies (21, 34) support the model depicted in **Figure 3D**. That is, *P. aeruginosa* overexpresses redox-active phenazines in co-culture with *Streptococcus* spp. which likely results in higher levels of H_2_O_2_, and both PerR and Dpr are important for *S. sanguinis* survival in this interaction.

## DISCUSSION

Building on previous knowledge of streptococcal interactions, we used a clinically-informed model community that simulates the CF lung environment to identify an important mechanism for *S. sanguinis* survival in the context of a CF model of polymicrobial infection. We showed that at a 24h timepoint, the *dpr* gene, encoding a DNA-binding ferritin-like protein, is robustly down-regulated in the mixed community setting but, when mutated, causes a defect in *S. sanguinis* SK36 viability in the mixed community (**Figure 1**). The *dpr* gene shows relatively high expression in monoculture, yet, when mutated, shows no growth defect compared to the WT when grown in monoculture using the CF-like conditions of ASM under anaerobic conditions. It is important to note that our transcriptional study only includes data collected at 24 hours post-incubation (39), while our time course data shown in **Figure 2B** suggests that *dpr* expression is important for *S. sanguinis* survival in the first 12 hours of growth. Furthermore, transcriptional data from the same study suggests that *P. aeruginosa* and *S. aureus* also apparently respond to oxidative stress in monoculture compared to the mixed community. Notably, in *S. aureus*, a thioredoxin reductase, *trxA*, and superoxide dismutase, *sodM*, are highly expressed in the mixed community compared to the monoculture. Furthermore, our transcriptional data indicate that for *S. sanguinis* SK36 there may be a reduction in ROS exposure, but even this reduced amount can be toxic to a cell lacking PerR or Dpr.

Here, we have begun to explore the mechanism(s) whereby the polymicrobial community causes a viability defect in the Ssx_*dpr* and Ssx_*perR* mutants. Based on previous work (34), we hypothesized that *P. aeruginosa* secreted, redox-active phenazines might be a contributor to the observed phenotype. Our data support this hypothesis, as blocking pyocyanin production via mutation of the *phz* genes successfully rescues the reduced growth phenotype of the Ssx_*dpr* and Ssx_*perR* mutants (**Figure 3A**). Similarly, exposing these mutants to the commercially available phenazine pyocyanin also reduced their viability compared to the WT. Consistent with our findings here, we showed previously that growth of a *P. aeruginosa* PA14 mutant in this same mixed community system produced phenazines ((25).

Our data also show that the addition of H_2_O_2_ can cause a reduction in the viability of the Ssx_*dpr* and Ssx_*perR* mutants (**Figure 3C**). Previous findings demonstrated that *P. aeruginosa* strains that overproduce pyocyanin display enhanced H_2_O_2_generation, cell lysis, and eDNA release, while strains that are unable to produce pyocyanin generate negligible amounts of H_2_O_2_ and released less eDNA (34). It is thought that pyocyanin promotes eDNA release in *P. aeruginosa* by inducing cell lysis mediated via H_2_O_2_ production, and the eDNA in turn facilitates *P. aeruginosa* biofilm formation ((34, 40–42). The correlation between increased phenazine production and elevated H_2_O_2_ levels in *P. aeruginosa* provided a rationale for the observed phenotype of the *S. sanguinis* Ssx_*dpr* and Ssx_*perR* mutants. That is, it may not be phenazine toxicity directly but rather phenazine-induced H_2_O_2_ production that require PerR-or Dpr-mediated protection. Our data demonstrate that both PerR and Dpr play an important role in protecting *S. sanguinis*, a catalase-negative bacteria, from H_2_O_2_.

Previous reports show that Fur-family transcriptional regulators PerR and OxyR can act as positive or negative regulators of oxidative stress response genes, including *dpr* and *sodA* (31, 32, 43–47). Our data shows that the Ssx_*perR* mutant phenotype in the mixed community mirrors that of the Ssx_*dpr* mutant (**Figure 2**). Therefore, we hypothesized that the observed viability reduction in the mixed community for both of these mutants relies on PerR regulation of *dpr*. However, our gene expression data suggests that PerR is a negative regulator of *dpr* under these conditions (**Figure 2C**). These findings indicate that the loss of viability in the Ssx_*perR* mutant may not be caused by a reduction in *dpr* expression. Interestingly, our findings also show that *sodA* expression is decreased in the Ssx_*perR* mutant, indicating that PerR is a positive regulator of *sodA* under these conditions (**Figure 2C**). However, a Ssx_*sodA* mutant does not show a reduction in viability in the mixed community (**Figure 1C**). Taken together, these data suggest that the loss of viability in the Ssx_*perR* mutant could result from a reduction in expression of a combination of PerR-induced genes. Alternatively, it is possible that the increased expression of *dpr* in the Ssx_*perR* mutant contributes to the observed Ssx_*perR* mutant phenotype by an unknown mechanism. Altogether, our findings suggest that further investigation of the PerR regulon is necessary to understand the complex inter-microbe dynamics involving streptococci in the CF airway.

Our results also suggest that phenazine and/or H_2_O_2_ production could be an antagonistic tactic employed by *P. aeruginosa* to compete with streptococci and other microbes in the CF airway. Dps-family proteins, the structure and function of which have been extensively studied across many genera of bacteria, but especially *E. coli*, have been shown to provide multifaceted protection to cells through DNA binding, iron sequestration, and its ferroxidase activity (48–53). Specifically in streptococci, previous studies have shown that strains without functioning Dps-family proteins, such as Dpr, display hypersensitivity to H_2_O_2_. Protection from H_2_O_2_ is mediated through the Dpr proteins iron binding activity (44, 54, 55). Other genetic factors have also been shown to contribute to H_2_O_2_ protection in streptococci, such as thiol peroxidases, superoxide dismutases, and NADH oxidases ((33, 56, 57). However, our data showing a lack of a phenotype for the thiol peroxidase or the superoxide dismutase mutants of *S. sanguinis* (**Figure 1**) suggests that PerR or Dpr-mediated H_2_O_2_ resistance are the most important mechanisms of ROS protection for *S. sanguinis* SK36 in our clinically relevant model.

Interestingly, our findings here complement previous studies showing that *Streptococcus* spp. can also negatively impact *P. aeruginosa* growth and viability via production of ROS and reactive nitrogen species (RNS) (17, 18). It is important to note that these previous studies were done in Todd-Hewit broth (THB) under aerobic growth conditions; thus, differences in the observed microbial interaction among these studies may be attributable to differential growth conditions.

In conclusion, this research contributes to our understanding of the intricate microbial interactions within the CF lung microbiome, emphasizing the role of *S. sanguinis* SK36 ROS response in the presence of *P. aeruginosa*. Previous findings indicating that lung microbiome composition can have significant effects on clinical outcome for pwCF (6, 8–10) underscoring the need for a better understanding of microbial dynamics for the development of effective interventions in the challenging landscape of CF lung disease. There are clinical data implicating that high *Streptococcus* abundance can lead to varying clinical outcomes, some positive and some negative for pwCF (4, 5, 11–15). Gaining a deeper understanding of streptococci dynamics in the CF lung could shed light on why the current data presents such contradictory results. Finally, we argue that continued study of microbe-microbe interactions, and further exploration into the broader implications of these interactions and their clinical relevance, is necessary to advance the capacity to manage and effectively treat the chronic infections associated with cystic fibrosis and other chronic airway infections.

## MATERIALS AND METHODS

### Bacterial strains and culture conditions

*P. aeruginosa* PA14*, S. aureus* Newman*, S. sanguinis* SK36, and *P. melaninogenica* ATCC 25845 strains used in this study are listed in **Supplemental Table S2**. All strains were cultivated as previously described (25) *P. aeruginosa* and *S. aureus* cultures were grown in tryptic soy broth (TSB) with shaking at 37°C. *P. melaninogenica* cultures were grown in TSB supplemented with 0.5% yeast extract (YE), 5 µg/mL hemin, 2.85 mM L-cysteine hydrochloride, and 1 µg/mL menadione (*Prevotella* growth medium [PGM]). *S. sanguinis* overnight cultures were grown in Todd-Hewitt broth supplemented with 0.5% YE (THY) at 37°C with 5% CO_2_. Artificial sputum medium with 5mg/mL mucin (ASM) was prepared as previously reported (25) and supplemented with 100 mM 3-morpholinopropane-1-sulfonic acid (MOPS) adjusted to a pH of 6.80.

### Complementation constructs for *S. sanguinis* mutants

For complementation of each targeted *S. sanguinis* gene, the suicide vector pJFP126 was used to allow for the insertion of complementing genes into an ectopic chromosomal site (*SSA_0169*) via homologous recombination and expression of each gene is under the control of an IPTG inducible promoter *hyper-spank* as previously reported (58). Primers and plasmids are reported in **Supplemental Table S2**.

### Statistical analysis

Data visualization and statistical analysis were performed using GraphPad Prism 9. The statistical test used and the resulting P value are listed in the appropriate figure legend or in the text.

### Microbial growth assays

Culture experiments were performed as previously described by (25) with modifications. Cells from overnight liquid cultures of each strain being used were individually collected and washed twice (for *P. aeruginosa* and *S. aureus*) or once (for *S. sanguinis* and *P. melaninogenica*) in sterile phosphate-buffered saline (PBS) by centrifuging at 10,000 × *g* for 2 minutes. After the final wash, cells were resuspended in the ASM base (without mucin). The optical density (OD_600_) was then measured for each bacterial suspension and diluted to an OD_600_ of 0.2 in ASM. Monocultures and co-cultures were prepared from the OD_600_ = 0.2 suspensions and further diluted to a final OD_600_ of 0.01 for each microbe in ASM. A polystyrene flat-bottom 96-well plate was inoculated with 100 µL of bacterial suspension for each monoculture and co-culture condition in triplicate. Plates were incubated using an AnaeroPak-Anaerobic container with a GasPak sachet (ThermoFisher) at 37°C for 24 hours. After incubation, unattached cells were aspirated with a multichannel pipette. Pre-formed biofilms were then washed once with sterile PBS and then resuspended in 50 µL PBS using a sterile 96-pin replicator. Each culture was 10-fold serially diluted and plated on selective media to quantify colony forming units (CFUs), as previously reported (25).

### Time course assays

Cultures for time course assays were prepared as described in the ‘Microbial growth assays’ section above. A separate 96-well plate was inoculated for each time point to be measured. At 6, 12, 18 and 24 hours, biofilms from one 96-well plate were collected and plated to quantify CFUs as described in ‘Microbial growth assays.’ Experiments were performed in triplicate on different days.

### qPCR assay

For qPCR experiments, *S. sanguinis* monocultures were cultivated as described above in the ‘Microbial growth assays’ section. At 24 hours, bacterial lysis and RNA extraction were performed following the protocols outlined in the RNeasy Mini Kit (Qiagen). 20μL mutanolysin (Sigma-Aldrich) was added to the lysozyme solution to aid cell lysis. RNA was treated with DNAse using the Ambion TURBO DNA-free Kit (ThermoFisher) and quantified by Nanodrop. 500ng of total RNA was reverse-transcribed using the RevertAid First Strand cDNA Synthesis Kit (ThermoFisher). Gene expression was quantified using the 2X iQ SYBR Green supermix (Bio-Rad). All primers used for qPCR are listed in **Supplemental Table S2**. Reactions containing non-reverse-transcribed RNA and no template were used as controls. We used transcriptional data from our lab’s previously reported RNA-Seq analysis to select a reference gene that shows minimal differential expression across conditions in a CF-relevant polymicrobial model system (28). *ntpA*, a V-type sodium ATPase catalytic subunit A, was selected from these data and used to normalize RNA samples. Relative gene expression was calculated using the ΔΔ*C*_T_ method.

### Phenazine and ROS response assays

A 20 mM PYO (Sigma) stock and a 20 mM PCA (Sigma) stock were each prepared in 100% DMSO. We used commercially available 30% stock solution of H_2_O_2_ (Fisher). To test phenazine and ROS response, *S. sanguinis* monocultures were grown as described in ‘Microbial growth assays’ for 24 hours. After the 24-hour incubation, unattached cells were aspirated with a multichannel pipette and discarded. 100μL ASM supplemented with 50uM, 150uM or 300uM of the phenazines pyocyanin or phenazine-carboxilic acid (PCA) or 0.1%, 0.2%, and 0.3% H_2_O_2_, depending on the condition, was added to wells, in triplicate. ASM supplemented with 0.5% DMSO was used as a control. The cultures were then incubated for an additional 24 hours in anoxic conditions at 37°C. After 48 total hours of incubation, biofilm CFUs were quantified as described in the ‘Microbial growth assays’ section. Experiments were performed in triplicate on different days.

### Supernatant assays

To test effects of supernatant, *S. sanguinis* monocultures were prepared as described in ‘Microbial growth assays’ with modifications. The OD_600_ of 0.2 for bacterial suspensions were diluted into medium consisting of 50% ASM and 50% prepared supernatant. Biofilm CFUs were quantified at 24 hours as described in the ‘Microbial growth assays’ section. Experiments were performed in triplicate on different days.

Supernatant was prepared by growing a full 96-well plate containing either a mix of *P. aeruginosa, S. aureus, S. sanguinis*, and *P. melananogenica*, or a monoculture of *P. aeruginosa*, depending on the experimental condition. These cultures were prepared as described in ‘Microbial growth assays.’ A 96-well plate containing uninoculated ASM was prepared as described below to use as a ‘no supernatant’ control. After a 24-hour incubation, the unattached cultures were aspirated with a multichannel pipette and transferred to sterile 2.0 mL microtube. Samples were then centrifuged at 16,000 *x g* for 10 minutes. The liquid portion was then removed from the microtube and passed through a 0.22 μm filter. The prepared cell-free supernatant was immediately added to ASM and inoculated per the methods described above. Sterility of the prepared supernatants was confirmed by spotting 5μL of the supernatant onto two TSB and two blood agar plates that were then incubated both aerobically and anaerobically at 37°C for 24 hours.

## ACKNOWLEDGEMENTS

This work was supported by National Institutes of Health (R01 AI155424) to G.A.O.

## Notes

### Competing Interest Statement

The authors have declared no competing interest.

### Summary of Updates

We noted an issue with an RNA-Seq file with corrupted data that reversed the fold change values, requiring us to revise Figure 1A, Supplemental Figure S1, and Supplemental Table S1. None of the other data were impacted by this issue. We also modified/updated a few areas in the text (results, discussion) to align with the revised data.

