## Supplemental Figures for "DPR-MEDIATED H_2_O_2_ RESISTANCE CONTRIBUTES TO STREPTOCOCCI SURVIVAL IN A CYSTIC FIBROSIS AIRWAY MODEL SYSTEM"

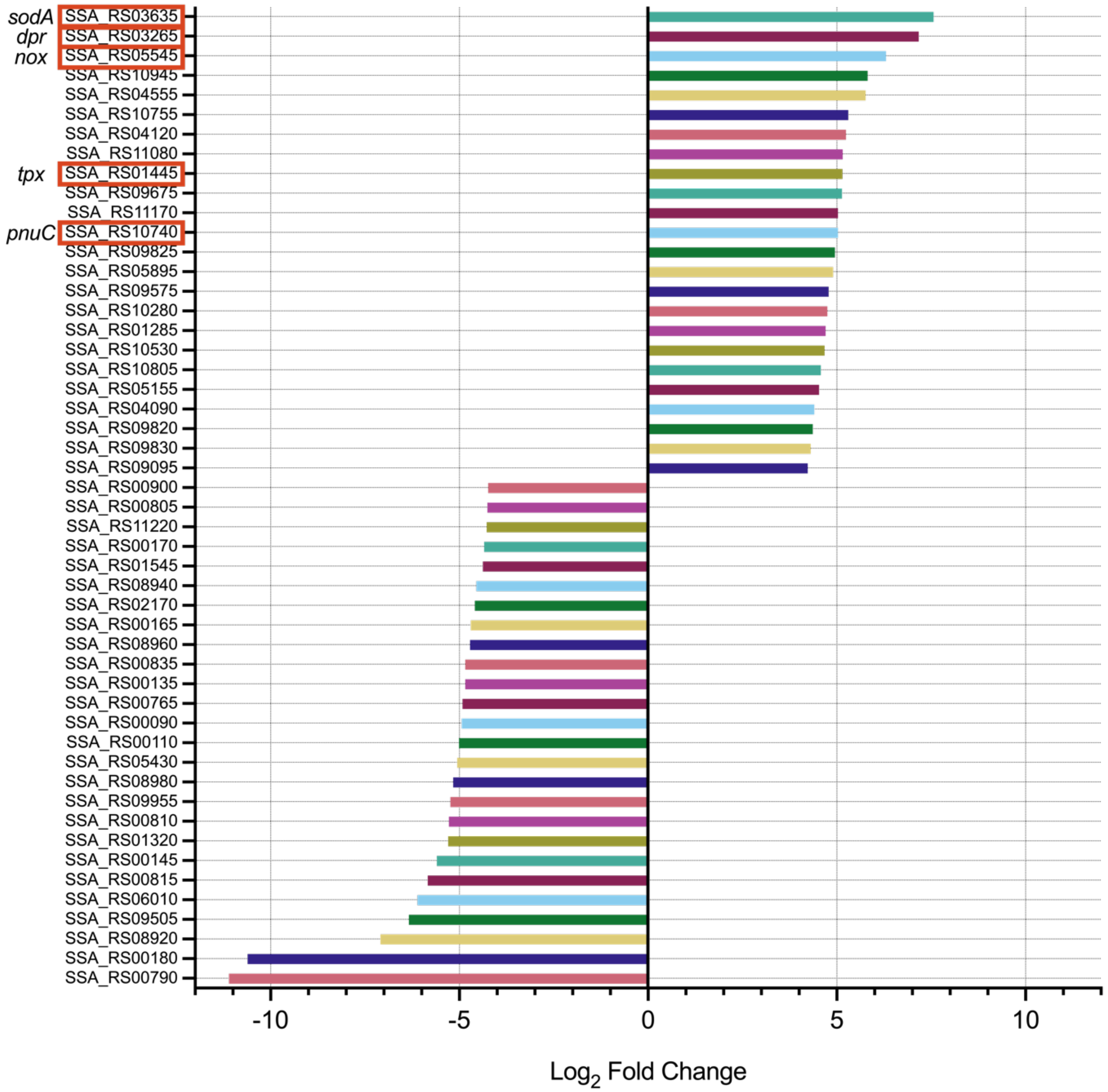


**Figure S1. *S. sanguinis* genes differentially expressed in the monoculture compared to the mixed community.** $Log_2$ fold change for the top fifty most differentially expressed *S. sanguinis* genes in the monoculture compared to the mixed community (genes up-regulated in the monoculture are positive). Genes selected for this study are boxed in orange. Differential expression was calculated using the DESeq2 package in R as previously reported (1).

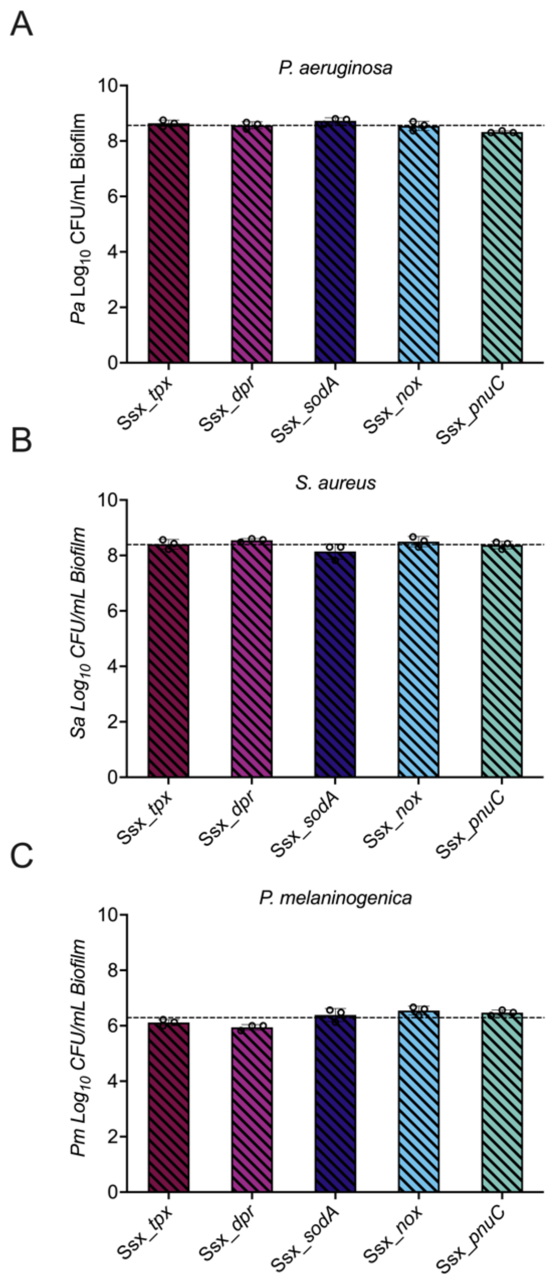


**Figure S2. The Ssx_*tpx,* Ssx_*dpr*, Ssx_*sodA,* Ssx_*nox,* and Ssx_*pnuC* mutants do not impact growth of other microbes in the mixed community.** (**A**) CFU counts for *P. aeruginosa* grown in the mixed community with the *S. sanguinis* mutants listed. Dashed line shows average CFU count for *P. aeruginosa* grown in the mixed community with WT *S. sanguinis*. CFUs were performed by plating on Difco *Pseudomonas* Isolation Agar (PIA). (**B**) CFU counts for *S. aureus* grown in the mixed community with the *S. sanguinis* mutants listed. Dashed line shows average CFU count for *S. aureus* grown in the mixed community with WT *S. sanguinis*. CFUs were performed by plating on Difco Mannitol Salt Agar (MSA). (**C**) CFU counts for *P. melaninogenica* grown in the mixed community with the *S. sanguinis* mutants listed. Dashed line shows average CFU count for *P. aeruginosa* grown in the mixed community with WT *S. sanguinis*. CFUs were performed by plating on medium selective for the growth of *Prevotella* spp. Each data point presented in a column represents the average from at least three technical replicates performed on at least three different days (n=3). Statistical analysis was performed using ordinary one-way ANOVA and Tukey's multiple comparisons posttest with *, $p<0.05$; ****, $p<0.0001$, no notation = non-significant. Error bars represent SD.

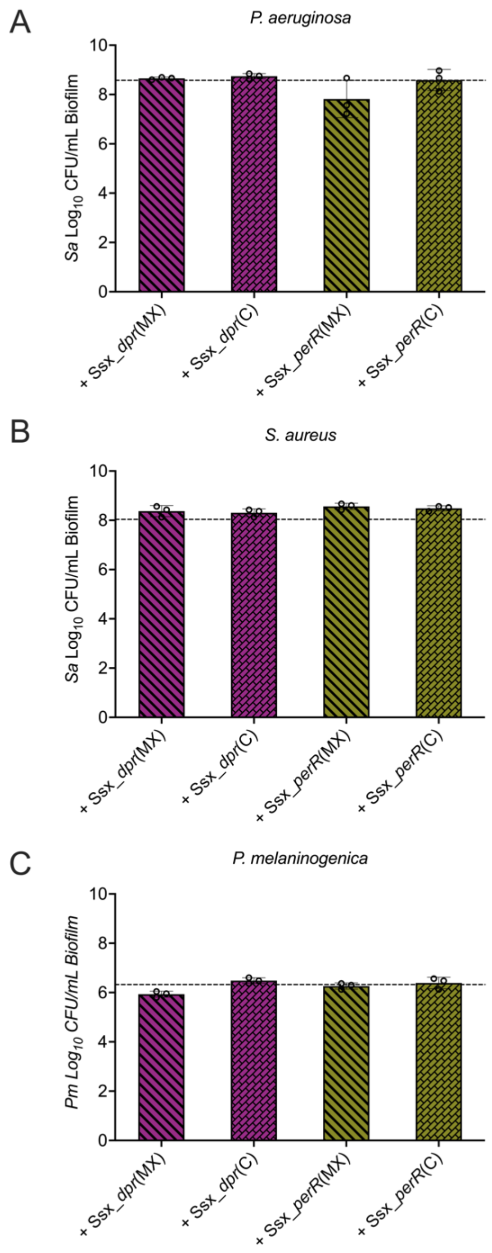


**Figure S3. The Ssx_*perR* mutant has little impact on the growth of other microbes in the mixed community.** (**A**) CFU counts for *P. aeruginosa* grown in the mixed community with *S. sanguinis* Ssx_*dpr* and Ssx_*perR* mutants and their complements. Dashed line shows average CFU count for *P. aeruginosa* grown in the mixed community with WT *S. sanguinis*. CFUs were performed by plating on Difco PIA. (**B**) CFU counts for *S. aureus* grown in the mixed community with *S. sanguinis* Ssx_*dpr* and Ssx_*perR* mutants and their complements. Dashed line shows average CFU count for *S. aureus* grown in the mixed community with WT *S. sanguinis.* CFUs were performed by plating on Difco MSA. (**C**) CFU counts for *P. melaninogenica* grown in the mixed community with *S. sanguinis* Ssx_*dpr* and Ssx_*perR* mutants and their complements. Dashed line shows average CFU count for *P. melaninogenica* grown in the mixed community with WT *S. sanguinis*. CFUs were performed by plating on medium selective for the growth of *Prevotella* spp. Each data point presented in a column represents the average from at least three technical replicates performed on at least three different days (n=3). Statistical analysis was performed using ordinary one-way ANOVA and Tukey's multiple comparisons posttest with *, p<0.05; ****, p<0.0001, no notation = non-significant. Error bars represent SD.

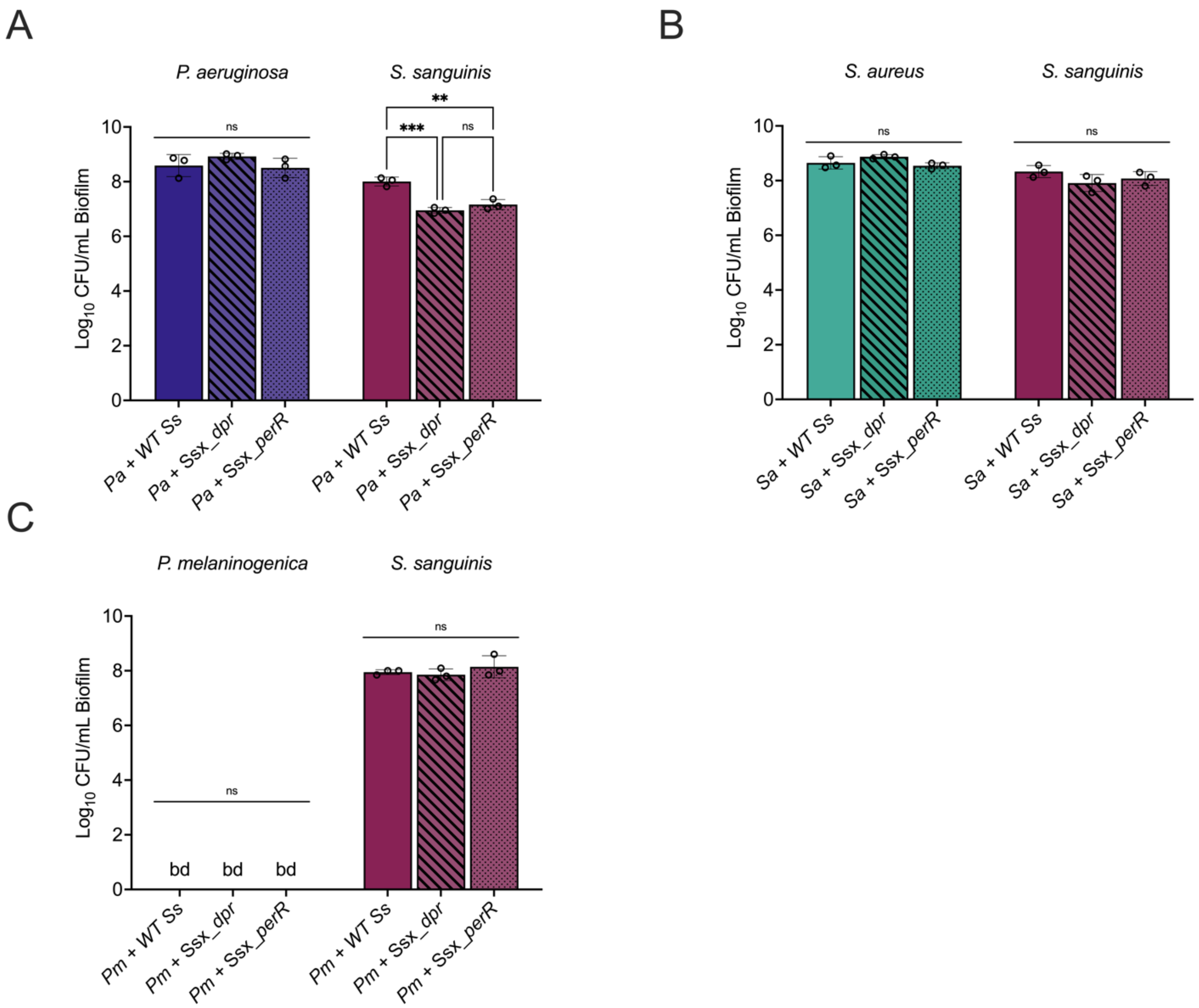


**Figure S4. Pairwise co-culture with *P. aeruginosa*, but not *S. aureus* or *P. melaninogenica*, replicates the *S. sanguinis* Ssx_*dpr* and Ssx_*perR* mutant phenotype observed in the mixed community.** (**A**) CFU counts for *P. aeruginosa* and listed *S. sanguinis* strains grown in co-culture. CFUs were performed by plating on Difco PIA and medium selective for *Streptococcus* spp. (**B**) CFU counts for *S. aureus* and listed *S. sanguinis* strains grown in co-culture. CFUs were performed by plating on Difco MSA and medium selective for *Streptococcus* spp. (**C**) CFU counts for *P. melaninogenica* and listed *S. sanguinis* strains grown in co-culture. CFUs were performed by plating on medium selective for *Prevotella* spp. or *Streptococcus* spp. Each data point presented in a column represents the average from at least three technical replicates performed on at least three different days (n=3). Statistical analysis was performed using ordinary one-way ANOVA and Tukey's multiple comparisons posttest with *, $p<0.05$; ****, $p<0.0001$, ns = non-significant. Error bars represent SD.

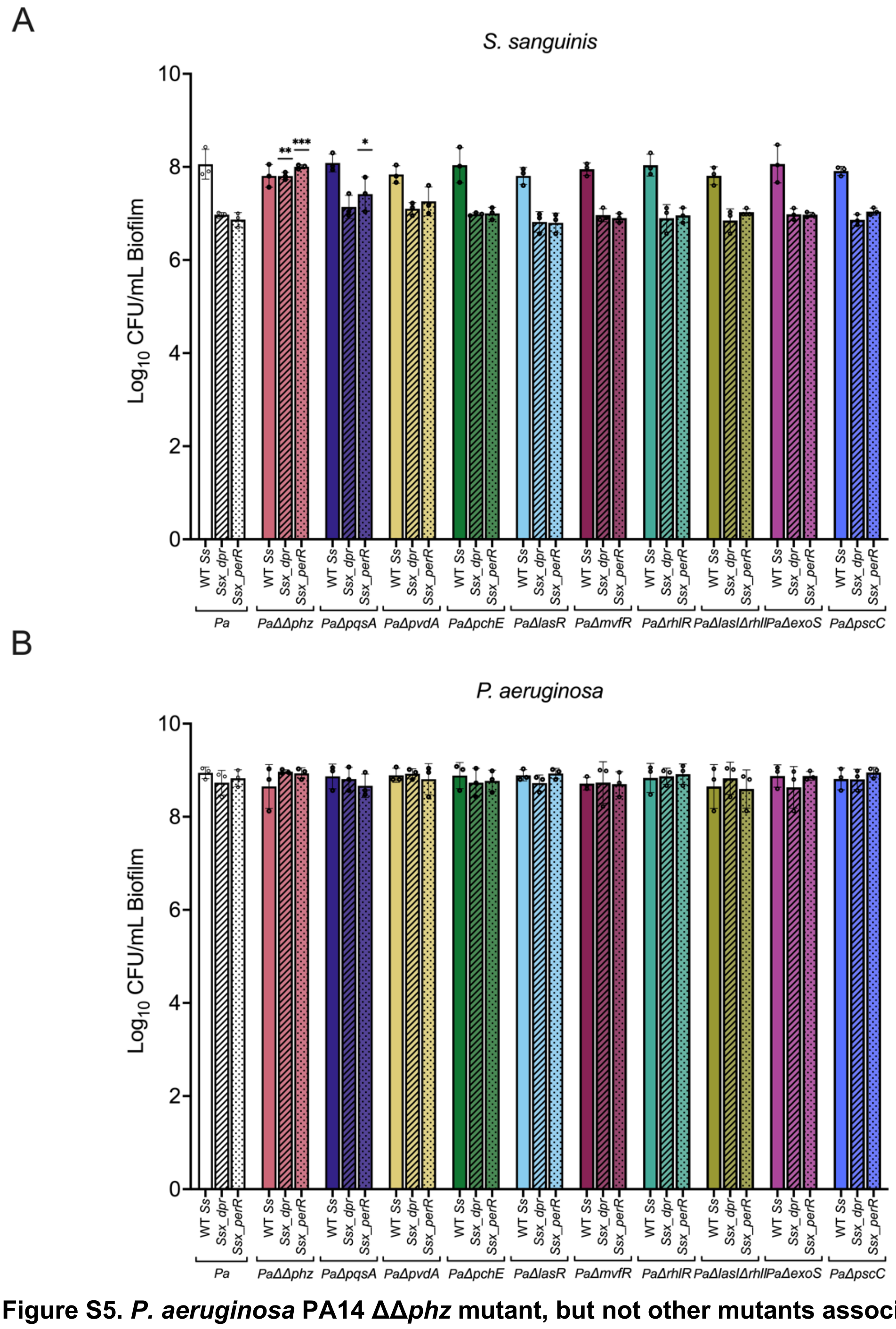


**Figure S5. *P. aeruginosa* PA14 ΔΔ*phz* mutant, but not other mutants associated with *P. aeruginosa* virulence, fully rescues the *S. sanguinis* Ssx_*dpr* and Ssx_*perR* viability defect observed in the co-culture.** (**A**) CFU counts for *S. sanguinis* WT, Ssx_*dpr,* and Ssx_*perR* grown in co-culture with *P. aeruginosa* mutants listed. CFUs were performed by plating on medium selective for *Streptococcus* spp. (**B**) CFU counts for listed *P. aeruginosa* mutants grown in co-culture with *S. sanguinis* WT, Ssx_*dpr,* and Ssx_*perR*. CFUs were performed by plating on Difco PIA. Each data point presented in a column represents the average from at least three technical replicates performed on at least three different days (n=3). Statistical analysis was performed using ordinary one-way ANOVA and Tukey's multiple comparisons posttest with *, $p<0.05$; ****, $p<0.0001$, no notation = non-significant. Error bars represent SD.

**Literature Cited**.

1. Kesthely CA, Rogers RR, Hafi BE, Jean-Pierre F, O'Toole GA. 2023. Transcriptional profiling and genetic analysis of a cystic fibrosis airway-relevant model shows asymmetric responses to growth in a polymicrobial community. *Microbiol Spectr* 11:201-223.
