## Supplemental Table S2 for "DPR-MEDIATED H_2_O_2_ RESISTANCE CONTRIBUTES TO STREPTOCOCCI SURVIVAL IN A CYSTIC FIBROSIS AIRWAY MODEL SYSTEM"

**Table S1. Strains, plasmids and primers used in this study.**

| STRAIN | SMC # | GENOTYPE | SOURCE |
|---|---|---|---|
| *Pseudomonas aeruginosa* PA14 | 232 | Rif(r) wildtype strain | (1) |
| PA14ΔΔ*phz* | 5020 | in-frame deletions of *phzA1-G1* and *phzA2-G2* genes | (2) |
| PA14Δ*pqsA* | 5013 | in-frame deletion of *pqsA* gene | (3) |
| PA14Δ*pvdA* | 9596 | in-frame deletion of *pvdA* gene | (4) |
| PA14Δ*pchE* | 6597 | in-frame deletion of *pchE* gene | (4) |
| PA14Δ*lasR* | 5021 | in-frame deletion of *lasR* gene | (5) |
| PA14Δ*mvfR* | 5018 | in-frame deletion of *mvfR* gene | (6) |
| PA14Δ*rhlR* | DH2742 | in-frame deletion of *rhlR* gene | (7) |
| PA14Δ*lasI*Δ*rhlI* | DH242 | in-frame deletion of *lasI* and *rhlI* genes | (8) |
| PA14Δ*exoS* | 1214 | in-frame deletion of *exoS* gene | D.A. Hogan |
| PA14Δ*pscC* | 2196 | in-frame deletion of *pscC* gene | (9) |
| *Staphylococcus aureus* Newman | 1007 | wildtype | (10) |
| *Streptococcus sanguinis* SK36 | 7474 | wildtype | (11) |
| Ssx_*tpx* | SK36-lib[a] | SK36 ΔSSA_0259::aphA-3; Km$^r$ | (12) |
| Ssx_*dpr* | SK36 lib | SK36 ΔSSA_0644::aphA-3; Km$^r$ | (12) |
| Ssx_*sodA* | SK36 lib | SK36 ΔSSA_0721::aphA-3; Km$^r$ | (12) |
| Ssx_*nox* | SK36 lib | SK36 ΔSSA_1127::aphA-3; Km$^r$ | (12) |
| Ssx_*pnuC* | SK36 lib | SK36 ΔSSA_2194::aphA-3; Km$^r$ | (12) |
| Ssx_*perR* | SK36 lib | SK36 ΔSSA_0686::aphA-3; Km$^r$ | (12) |
| Ssx_*dpr*::*dpr* | | Ssx_*dps* complement | This Study |
| Ssx_*perR*::*perR* | | Ssx_*fur* complement | This Study |
| *Prevotella melananogenica* ATCC 25845 | 6965 | DSM 7089 / ATCC 25845 | (13) |

[a]Mutant isolated from the *Streptococcus sanguinis* SK36 mutant library collection (12).

| PLASMID | DESCRIPTION | SOURCE |
|---|---|---|
| pJFP126 | suicide vector with *hyper-spank* promoter | (14) |

| PRIMERS | SEQUENCE | USE |
|---|---|---|
| dpr_comp_F | TTTGATTGTTCCTCATAAAGATGGT | complementing Ssx_*dpr* |
| dpr_comp_R | GCCATTTATTCCTCCTAGTTAGTCAC | complementing Ssx_*dpr* |
| perR_comp_F | GTTTTAGTACCTGGAGGGAATAAT | complementing Ssx_*perR* |
| perR_comp_R | CGTTAGAACAAGTTTTTGCAGAGAT | complementing Ssx_*perR* |
| SSA_0091_RT_F | GGAGCGTTTCCAGACCATCA | real time PCR |
| SSA_0091_RT_R | CCGACAGACAGACCTGGAAC | real time PCR |
| dpr_RT_F | GGCTTCATGGTTTGGGATCC | real time PCR |
| dpr_RT_R | GGCTTGCCTCCCAGTGTAAT | real time PCR |
| perR_RT_F | GGCCATCAGCACCTCAATGT | real time PCR |
| perR_RT_R | AGTCGTTTGGCTTTGGGTGA | real time PCR |
| sodA_RT_F | CGCTTACGATGCTTTGGAGC | real time PCR |
| sodA_RT_R | GTTGATGACTGCCTGACGGA | real time PCR |

**Literature Cited.**
